# Cell-Cell Fusion in NSCLC Promotes Evolution of Therapeutic Resistance

**DOI:** 10.1101/2024.12.02.626399

**Authors:** Paulameena V. Shultes, Dagim S. Tadele, Arda Durmaz, Davis T. Weaver, Rowan Barker-Clarke, Mina N. Dinh, Siqing Liu, Endalkachew A. Alemu, Simon Rayner, Andriy Marusyk, Jacob G. Scott

## Abstract

Cell-cell fusion has been implicated in various physiological and pathological processes, including cancer progression. This study investigated the role of cell-cell fusion in non-small cell lung cancer (NSCLC), focusing on its contribution to chemoresistance and tumor evolution. By co-culturing drug-sensitive and drug-resistant NSCLC cell lines, we observed spontaneous cell-cell fusion events, particularly under gefitinib selection. These fused cells exhibited enhanced fitness and a higher degree of chemoresistance compared to parental lines across a panel of 12 chemotherapeutic agents. Further analysis, including fluorescence imaging and cell cycle analysis, confirmed nuclear fusion and increased DNA content in the fused cells. Bulk RNA sequencing revealed genomic heterogeneity in fused cells, including enrichment of gene sets associated with cell cycle progression and epithelial-mesenchymal transition, both of which are hallmarks of cancer. These findings demonstrate that cell-cell fusion can act as a novel source of chemotherapeutic resistance and further promote aggressive phenotypes in NSCLC, highlighting the potential of fusion as a therapeutic target.

## Introduction

Cell-cell fusion is a complex process characterized by cellular approach, adhesion, membrane pore opening, cytoplasmic mixing, and in some cases nuclear mixing.^1^ Despite this complexity, spontaneous cell-cell fusion occurs throughout the human body and contributes to key pathways in human physiology and pathology.^2^ Current literature suggests that in humans, the roles of cell-cell fusion vary from embryogenesis and placentation to promoting cancer malignant potential.^2–4^ There is increasing evidence of cell-cell fusion potentiating cancer malignancy, but the study of cell-cell fusion has not yet expanded to include non-small cell lung cancer (NSCLC).^2,4^

Non-small cell lung cancer accounts for the majority (∼ 85%) of lung cancers, and is one of the leading causes of cancer-related deaths worldwide.^5^ Genomic characterization has identified common oncogenes such as EGFR that serve as molecular targets for modern NSCLC chemotherapeutics.^5^ Current treatments for NSCLC vary in efficacy, and like many cancers, NSCLC patients can and often do evolve therapeutic resistance to the standard immunologic and chemotherapeutics through a variety of mechanisms, including immune evasion and Darwinian processes.^5–9^ A recent study identified that cancer cell-mesenchymal stem cell (MSC) fusion directly contributes to NSCLC immune evasion and decreased efficacy of PD-L1 immunologic agents.^10^ This study thus raises the question of how else cell-cell fusion can contribute to the increased rates of treatment failure in NSCLC patients.

In this study, we have isolated and characterized NSCLC (PC-9) cells after spontaneous cell fusion of drug-resistant and sensitive lineages to gefitinib, an EGFR inhibiting immunologic agent. We demonstrate the presence of cell-cell fusion with nuclear fusion through several mechanisms, including fluorescent imaging, fluorescent activated cell sorting (FACS) analysis, and quantification of DNA content and cell-cycle analysis. We then examine the potential of fused NSCLC cells to propagate therapeutic resistance across different types and classes of NSCLC treatment using a variety of collateral sensitivity screenings.

In addition, we analyze the genomic features that arise after fusion in NSCLC that could contribute to the malignant potential of NSCLC. We conclude that cell-cell fusion can serve to directly propagate and promote therapeutic resistance in NSCLC and that further study is needed to identify the full extent of the impact of cell-cell fusion on tumor evolution.

## Results

### Identification and Isolation of Cell-Cell Fusion in NSCLC

We identified cell-cell fusion in NSCLC during a long-term co-culture experiment designed to study the dynamics between drug-sensitive (parental) and drug-resistant (resistant) NSCLC cell populations.^11–13^ Parental and resistant cells were tagged with nuclear-localizing green fluorescent protein (GFP) and red fluorescing mCherry, respectively, and co-cultured at various initial ratios under control and gefitinib treatment conditions (20nM) for 12 weeks. Under control conditions, parental cells exhibited higher intrinsic fitness (as defined by per capita growth rate under no drug selection), resulting in an increased frequency, while resistant cells showed fitness costs and declined in frequency. In contrast, under gefitinib treatment, resistant cells gained a fitness advantage and predominated within the co-culture (**Figure 1A, B**).

**Figure 1.**
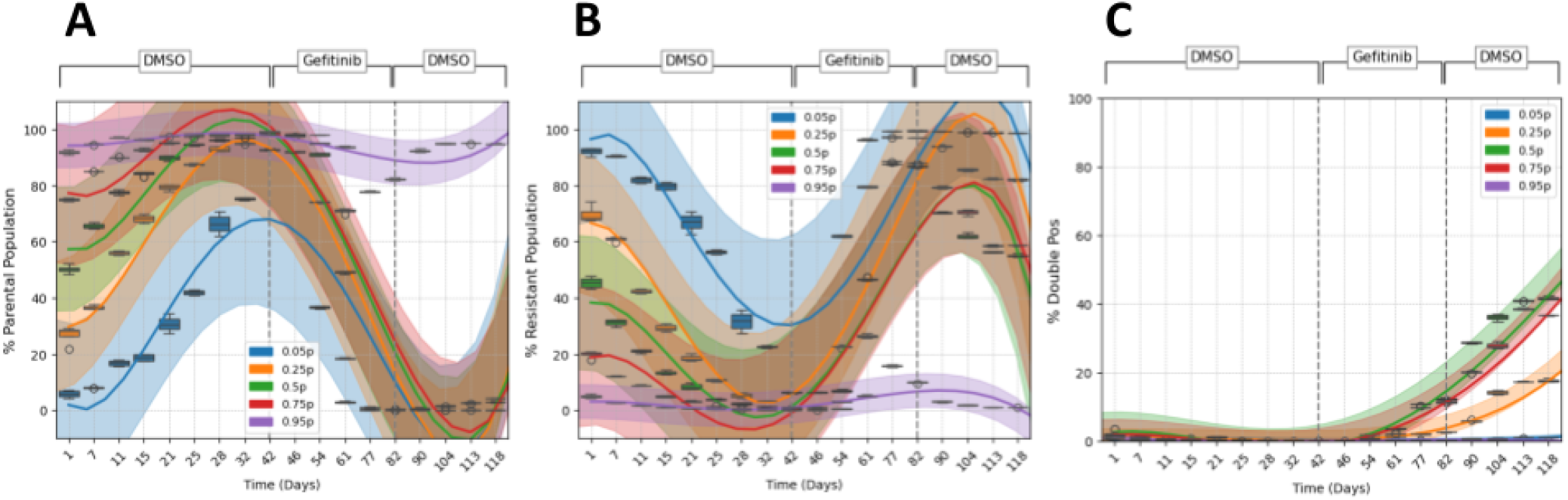
Repeated co-culture of sensitive (parental) and resistant cells at different ratios yielded different proportions of double-positive cells. ***A:*** In the absence of drug, the parental cell population reached 80-100% confluence across all co-culture ratios before dropping steadily under gefitinib selection. ***B:*** Resistant cells approach extinction in the absence of drug across co-culture ratios, and clonally expand rapidly following administration of gefitinib. ***C:*** Double-positive cells expressing both GFP (of parental cells) and mCherry (of resistant cells) occurred spontaneously at low (1-2%) rates prior to gefitinib selection based on cell-counting. Following prolonged selection, the double-positive cells reach varying levels of confluence 1-41% under different conditions. Co-culture of sensitive and resistant cells in a 50-50 split yielded the highest percentage of these double positive cells. Later study identified that the double-positive cells were derived from cell-cell fusion.

Remarkably, a third cell population, expressing both GFP and mCherry, emerged under gefitinib treatment. These yellow-fluorescing double-positive cells, which we later identified as being indicative of cell-cell fusion of mCherry and GFP fluorescing cells, exhibited enhanced fitness and underwent clonal expansion both in the presence of gefitinib and during a subsequent drug-free recovery period (treatment holiday) lasting six weeks. Double-positive cells were isolated via fluorescence-activated cell sorting (FACS) for further characterization.

Importantly, our quantitative analysis revealed that the double-positive cell population constituted over 40% of the total cells in co-cultures initiated with a 1:1 parental-to-resistant cell ratio. These levels can be attributed to a mixture of novel cell-cell fusion events and the replication/promotion of previously fused cells. Of interest was also the lack of double-positive population drop-off during the DMSO recovery periods. This suggests that there was some intrinsic fitness that supported double-positive persistence even as parental/resistant cell populations diminished. This data could also underlie unique ecological dynamics between parental, resistant, and double-positive cells that would require further study to parse. Double-positive cell counts were lower but still substantial in co-cultures with a 75:25 parental-to-resistant ratio (**Figure 1C**). Of note, the double-positive cells not only persisted during the treatment holiday, but continued to increase in cell count in contrast to their parental (GFP) and resistant (mCherry) counterparts. We hypothesize that this can be attributed to the effects of mixing the higher relative fitness of these cells (compared to parental and resistant cells in the absence of drug) and the complex ecological dynamics at play in this 3-player environment.^12^

Given that the fluorescent markers were nuclear-localizing, we theorized that these double-positive cells arose from cellular and nuclear fusion between the GFP-tagged parental and mCherry-tagged resistant cells. We further hypothesized this fusion would potentially allow for accumulation of novel variants and increased genomic heterogeneity promoting resistance. To monitor the frequencies of parental and resistant cell populations and detect potential fusion events, we employed high-parameter flow cytometry, which provides detailed insights into heterogeneous cell populations with high sensitivity. Flow cytometry analysis revealed a distinct double-positive population (over 41% of cells) expressing both GFP and mCherry. The size of the double-positive cell population far exceeded that which would likely occur from incidental doublet sorting. Thus, this sorting confirmed the probability of cell-cell fusion having occurred (**Figure 2**).

**Figure 2.**
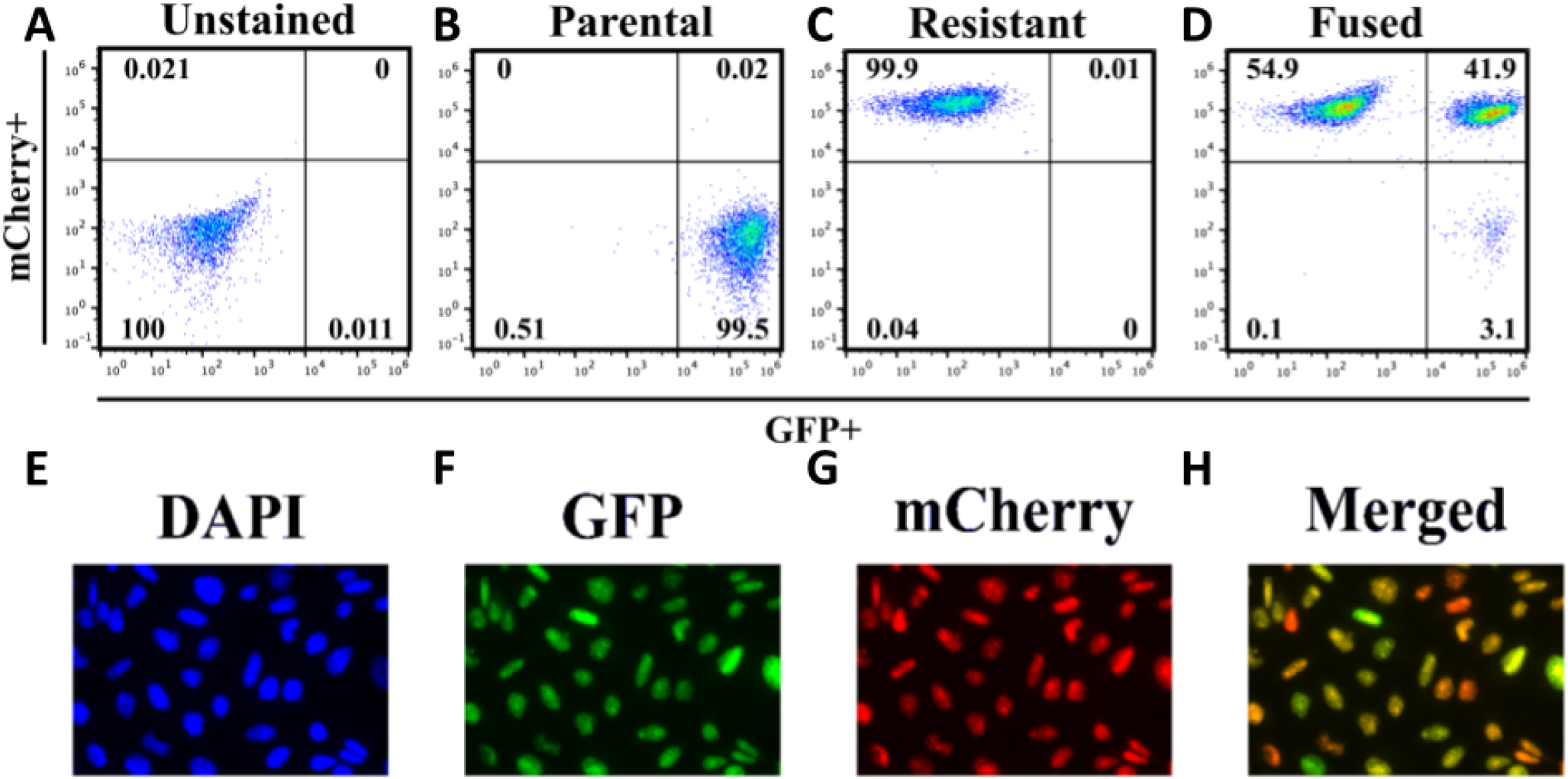
Fluorescent-activated cell sorting (FACS) isolates fused cells from cell-cell fusion with nuclear fusion following 50-50 co-culture experiments. **A:** Untagged PC 9 NSCLC cells served as a control, and demonstrated the absence of both GFP or mCherry signals at baseline using FACS. **B:** mCherry expression in isolated tagged resistant cell was observed in 99.85% of the cell population, confirming successful mCherry uptake. **C:** GFP expression in isolated tagged sensitive cells was observed in 99.88% of the cell population, confirming successful GFP uptake. **D:** Following long-term co-culture experiments with gefitinib selection, identification of over 41% fused cells was confirmed with 41.53% co-expression of GFP and mCherry signals following FACS. The remaining cells expressed either mCherry (55.11%) or GFP (3.34%) alone and represent remaining parental (GFP) and resistant(mCherry) cells in the population. **E:** DAPI staining of presumed fused cells isolates nuclei in fluorescence imaging. **F:** Parental PC 9 cells were tagged with nuclear-localizing GFP; imaging confirms retention of GFP fluorescence in isolated fused cells. **G:** Resistant PC 9 cells were tagged with nuclear-localizing mCherry; imaging confirms retention of mCherry fluorescence in isolated fused cells. **H:** Combined fluorescent images demonstrates presence of co-expression of mCherry and GFP in fused nuclei following cell-cell fusion. *FACS images of parental and resistant cell populations is available in Supplemental Figure 1*.

To confirm that the observed fusion events involved nuclear fusion specifically, we performed DAPI staining and fluorescence imaging to assess the co-localization of GFP and mCherry signals. The analysis revealed complete co-localization of GFP and mCherry fluorescence to the DAPI-stained nuclei of the double-positive cells (**Figure 2E**). This suggests not only that cell-cell fusion events occurred but that nuclear fusion occurred between the parental and resistant cells. These findings further highlight the significant potential of cell-cell fusion to promote genetic heterogeneity and mutagenesis in cancer.

To further explore and characterize the genotypic and phenotypic differences of our three cell populations, we proceeded with downstream analysis using the FACS-sorted double-positive cells (GFP + mCherry) and passage-matched parental (GFP) and resistant (mCherry) cell populations. From this point on, our double-positive cells will be referred to as fused cells. We hypothesized that the fused cells had undergone genetic (nuclear) mixing, so to confirm this hypothesis, we proceeded with a cell-cycle analysis to evaluate DNA content of the fused cells. In this experiment, we observed higher levels of DNA content in the fused cells across all phases of the cell cycle compared to either the parental or resistant cells alone **(Figure 3)**. On average, the DNA content in fused cells is not doubled, and this likely reflects the many potential pathways cells can take following cell-cell fusion to maintain or reduce their ploidy.^2–4,14^ Importantly, the higher-on-average DNA content suggests that the fused cells form a poly-aneuploid cell population, which has important implications for their relative fitness. Poly-aneuploidy cells have been repeatedly correlated with therapeutic resistance and stress tolerance within many biological contexts – varying from yeast populations, to human livers, and relevantly, human tumors.^15–19^ This serves as yet another hypothesis as to a potential mechanism by which cell-cell fusion in NSCLC could worsen clinical outcomes.

**Figure 3.**
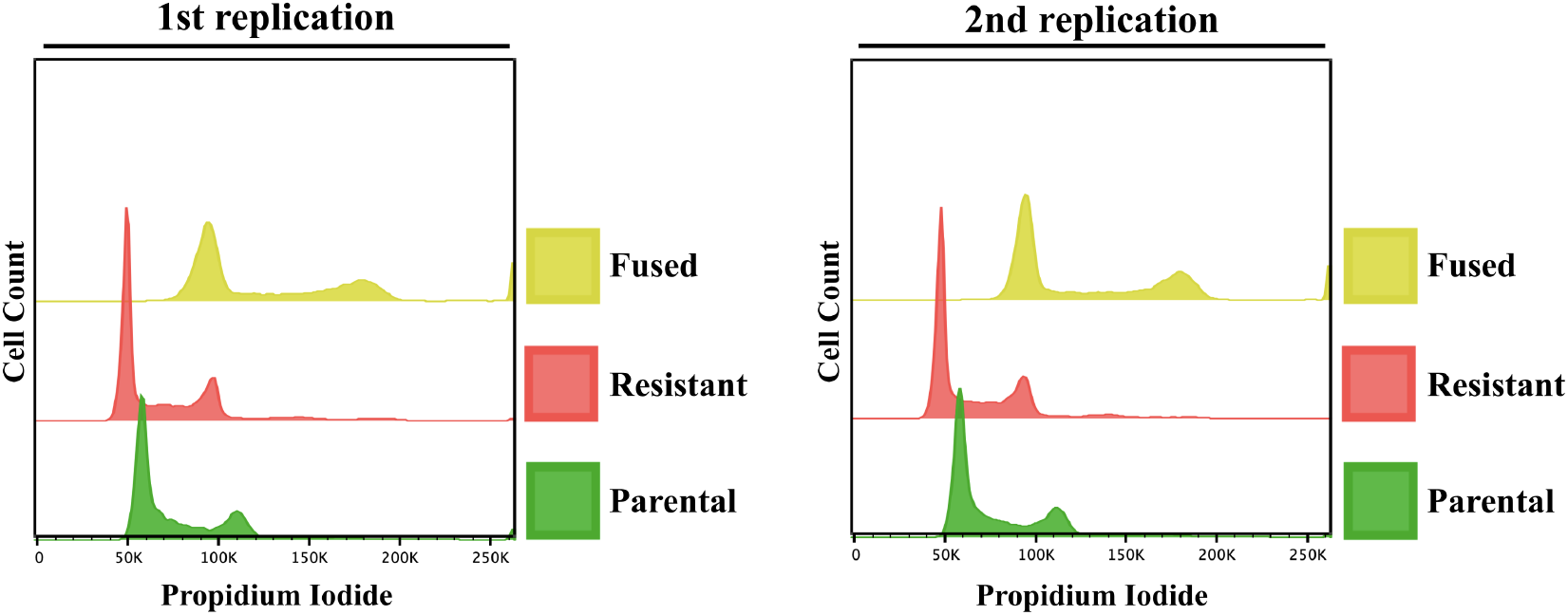
Cell-cycle analysis provides evidence of nuclear fusion by demonstrating higher DNA content of fused (Fused) cells relative to parental (Par) or resistant (Res) cells, across two experimental replicates. All three cell populations – parental, resistant, and fused – had similar cell counts in each phase of the cell cycle. The first spike represents G1, the lull in between the first and second spike represents S, and the second spike represents G2. The overall right-shift of the curve of the fused cells confirms higher than average levels of DNA content in each phase compared to the other two cell populations.

### Evidence of Conferred Fitness Advantages Secondary to Cell-Cell Fusion

Following our isolation of fused cells, we evaluated the role of cell-cell fusion in contributing to cancer malignant potential through two mechanisms: propagation of therapeutic resistance and promotion of cancer hallmarks.

#### Propagation of Drug Resistance

To characterize the effect of cell-cell fusion on therapeutic resistance in NSCLC cells, we performed a series of cell viability assays comparing the survival of our parental, resistant (evolved against gefitinib), and fused cell lines in the presence of 12 different therapeutic agents. When selecting therapeutic agents to test, we prioritized agents that are commonly used as first or second line therapies in the treatment of NSCLC. The drugs utilized spanned several different classes of treatment currently used to treat NSCLC, including EGFR inhibitors and other Tyrosine Kinase Inhibitors (TKIs), DNA repair inhibitors, and taxanes.^20,21^ Across the 12 drugs we evaluated, the fused cells consistently matched or exceeded the cell viability levels of the resistant cells at maximum drug concentrations, and both resistant and fused cells consistently outperformed the sensitive parental cells at those doses, as demonstrated in **Fig. 4a** Furthermore, as presented in **Fig. 4b**, IC50s were estimated following repeated bootstrapping via 4-parameter hill function where fused cells showed increased resistance compared to resistant cells for drugs erlotinib, gefitinib and pemetrexed. Conversly for trametinib, crizotinib and osimertinib, fused cells were in between the parental and resistant cells. Interestingly, for Cisplatin, Erlotinib and Trametinib we observed increased variance across bootstrap samples. (See supplemental figure 2 for individual dose-response curves) Fundamentally, both analyses demonstrate a generalized resistant phenotype among fused cells, but it is simultaneously clear that the level of selective pressure (drug dose) administered and tested for will lead to variability in specific measured outcomes. Deciding which of these metrics is most useful clinically requires further evaluation, but IC50 is more widely used when designing clinical interventions.

**Figure 4.**
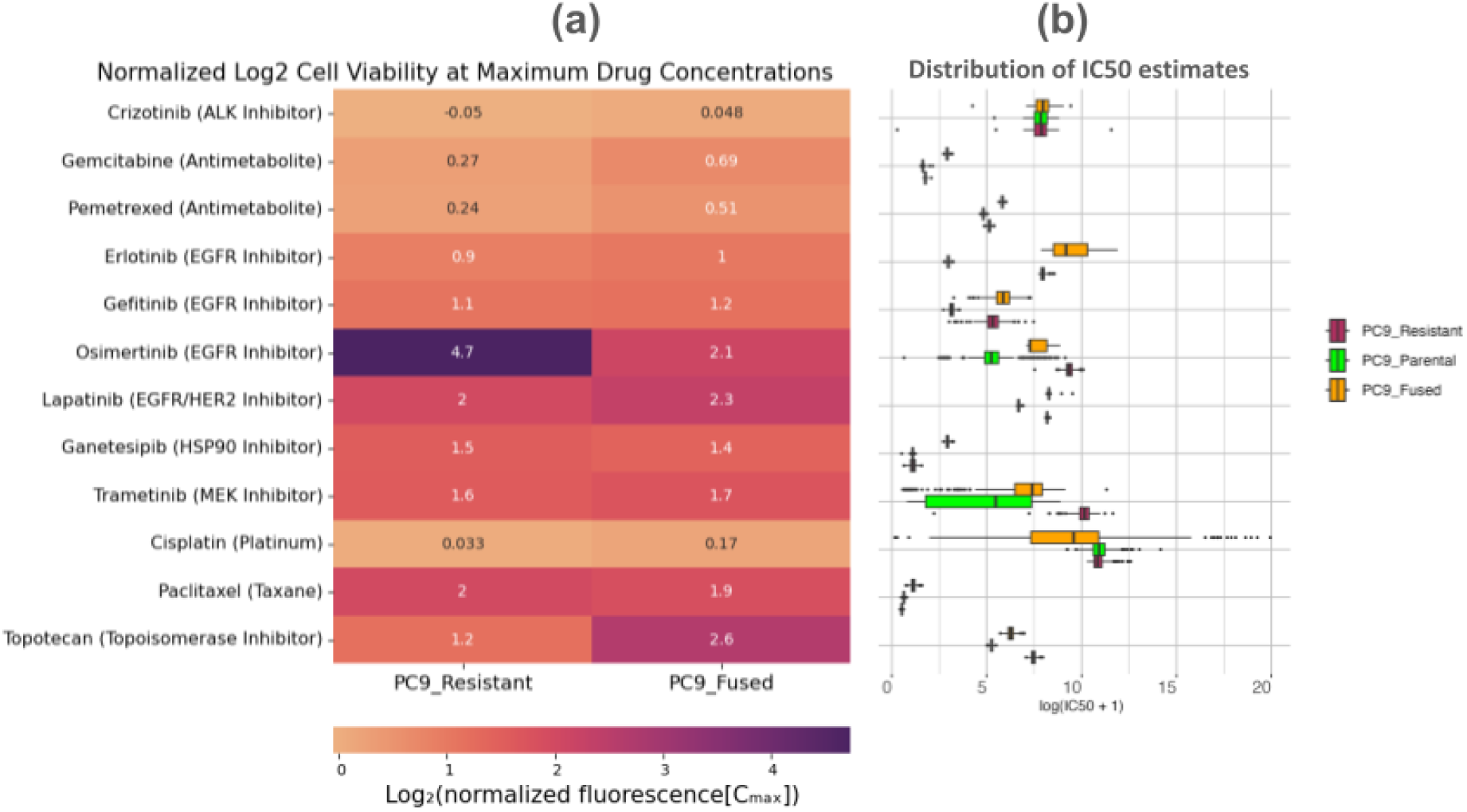
Fused cells illustrate a generalized resistant phenotype that often matches or exceeds that of the evolved-resistant cell line. **A:** When the cell viabilities of resistant and fused cell lines are normalized to the parental cell line at maximum drug concentrations, the fused cells persistently match or exceed the cell viability of the resistant cells. The key exception is osimertinib, which requires further evaluation. **B:** Distribution of bootstrap estimates of IC50 showing increased resistance of fused-cells across majority of the drugs tested. Panel B has a truncated X-axis for readability, full figure is available in supplements.

#### Bulk RNA-seq and Evidence of the Potentiation of Cancer Hallmarks

Previous studies have shown that spontaneous cell-cell fusion with nuclear fusion leads to increased genome instability and can potentiate tumor evolution in breast cancer.^4^ As another epithelial based cancer, we anticipated that NSCLC would have similar characteristics, especially following our observations of nuclear fusion. To evaluate the role of fusion in potentiating cancer hallmarks in NSCLC, we conducted differential gene expression and gene set enrichment analyses across our three cell lines of interest (parental, resistant, and fused). Given the range of changes observed in drug responses, we then investigated the transcriptional differences across samples. We restricted our analysis to genes with protein coding transcripts to reduce noise. Interestingly, top differentially expressed genes showed similar patterns between the resistant and fused samples whereby parental population were most distant from the fused population (**Figure 5a**). Subsequent gene set enrichment analysis revealed that the most differentially expressed genes in fused cells were centralized across several key processes including cell cycle regulation and epithelial mesenchymal transition (EMT) **Table 1** Given the directionality of change, we investigated further the EMT pathway which potentially is a transitioning state (**Figure 5b**). Fused cells showed decreased expression in certain genes compared to their resistant counterpart, yet still upregulated compared to parental (sensitive) cells. This is interesting given the strong correlation previous studies have drawn between cell-cell fusion events and the process of epithelial-mesenchymal transition.^2^ Upon closer inspection of **Table 1**, it becomes apparent that fused cells were in fact downregulated across all gene sets compared to resistant cells (refer to *fvr_NES* in **Table 1**). This surprising result is not what we had anticipated, as fusion has been phenotypically correlated with cancer hallmarks repeatedly in the literature.^2^ Further work is needed to understand how the fused cells can be so transcriptomically downregulated yet maintain a generally therapeutic resistant phenotype.

**Table 1.**
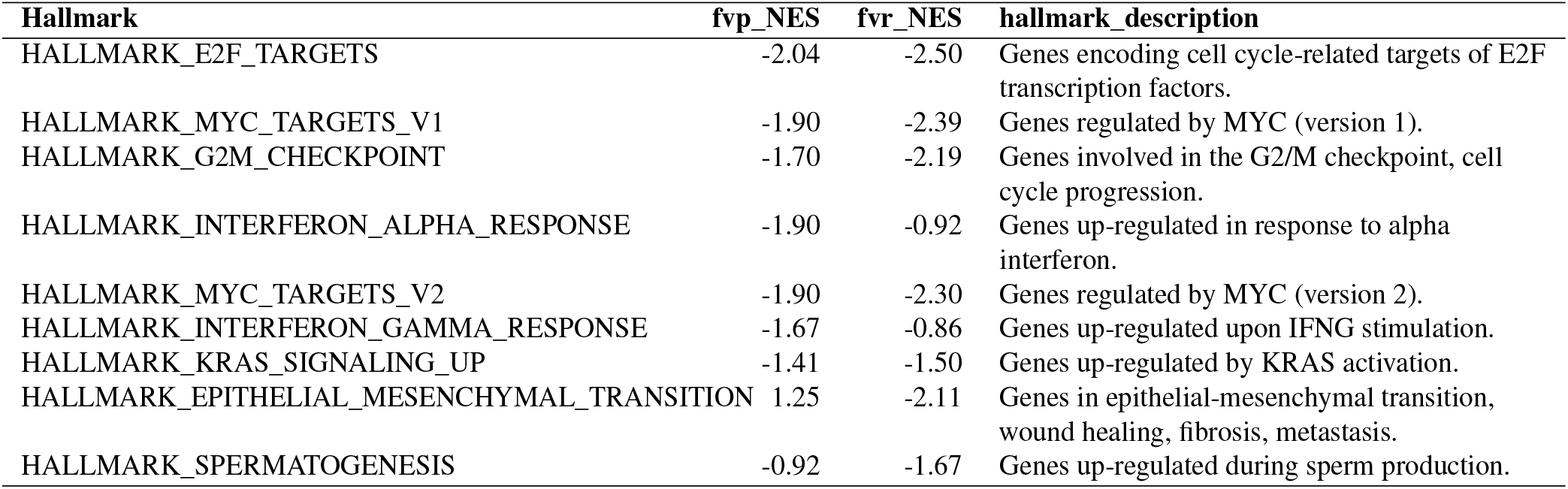
Gene set enrichment analysis (GSEA) of hallmark pathways with normalized enrichment scores (NES). **fvp_NES:** This represents the normalized enrichment score for each hallmark when comparing the fused cells versus parental cells. **fvr_NES:** Similarly, this represents the normalized enrichment score for each hallmark when comparing fused cells versus resistant cells. Importantly, fused cells demonstrate global down regulation across all hallmarks compared to both parental and resistant cells, with the exception of fused vs parental in the EMT pathway. A deeper dive into the EMT pathways is provided in **Figure 5b**

**Figure 5.**
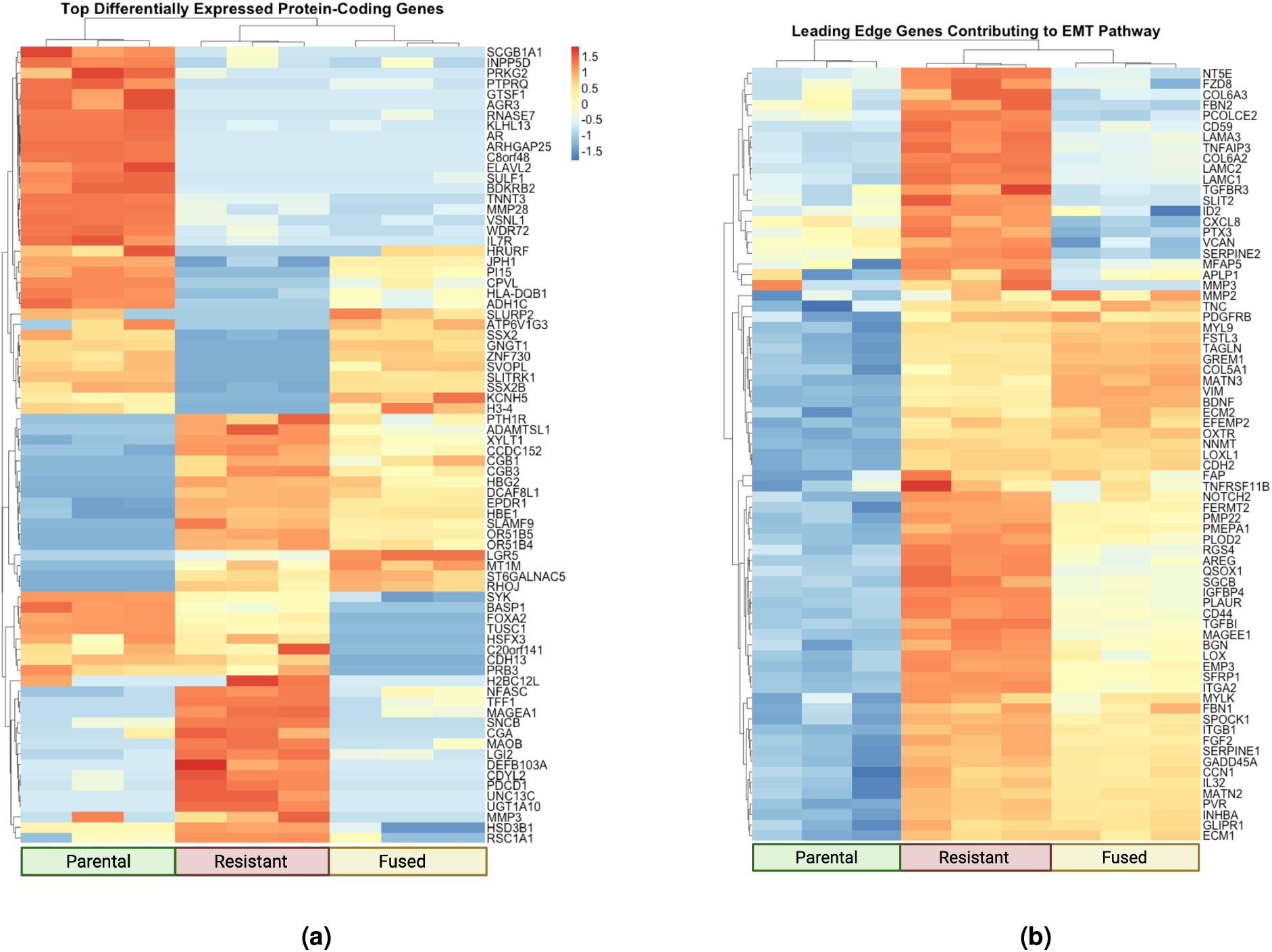
Comparison of transcriptomic heterogeneity in different bulk RNA-seq panels. **A:** Some genes seem to inherent the parental transcriptomic profile, while others inherent the resistant transcriptomic profile, and still others demonstrate new expression levels entirely. **B:** When the fused cells are compared to parental and resistant cells following pathway enrichment analyses, individual pathways demonstrate a similar level of mixed transcriptomic responses compared to the cell lines of origin. Some gene expression levels seem to reflect a transitionary state between the parental and resistant expression levels.

## Discussion

Cell-cell fusion is a complex process that occurs throughout human physiology and pathology.^2^ There has been increasing evidence that cell-cell fusion contributes to tumor malignancy through a variety of mechanisms related to the cancer hallmarks, from promoting cancer stemness and proliferative potential^22–25^ to enabling the epithelial-mesenchymal transition and metastasis.^25–27^ Despite these recent research advances, the process of cell-cell fusion has not been studied extensively in many cancers, including NSCLC.^2,4^ Prior studies have primarily investigated the role of cancer cell-somatic cell fusion in other epithelial cancers, with observed effects ranging from potentiating immune evasion^14,28–30^ and local invasion of surrounding tissues.^31^ Only recently have studies expanded their purview to consider intratumor cell-cell fusion and its potential to accelerate tumor evolution and rates of treatment failure.^4,32–34^

In this study, we isolated and characterized cells derived from the cell-cell fusion of two different EGFR-mut NSCLC (PC9) cell lineages, one with evolved resistance to gefitinib and one with sensitivity to gefitinib. Over the course of a prolonged co-culture experiment between our GFP-parental and mCherry-resistant PC-9 cells, we observed that cell-cell fusion occurred at constant low rates (1-2%) in the absence of drug. However, with treatment, selective pressure favored cells with resistant drug profiles including resistant and their fused counterparts. Furthermore, when collateral drug sensitivities were measured using cell viability assays, we observed strong evidence for collateral resistance. In fact, the improved cell viability of fused cells in high doses of chemotherapy was consistent across 12 different drugs used today in the control of NSCLC, across several different therapeutic classes and mechanisms of actions. The fused cells thus present with a general therapeutic resistance phenotype, which we initially hypothesized was inherited from their resistant cell lineage.

Subsequent to treatment characterization, we investigated the transcriptional profiles of fused cells in comparison to parental (PC-9) and resistant cells. Our preliminary analysis demonstrated a general downregulation in many key cellular processes and pathways following fusion between NSCLC cells, especially when compared to their resistant cell lineage. When looking at the key pathways such as EMT and cell cycle checkpoints, we see fused cells are not as transcriptionally active as either of their lineages (parental or resistant). This seems to suggest that the fused phenotype – including the general resistant phenotype – originates from processes separate from those that give the resistant cells their treatment resistance. Therefore, further study is needed to elucidate the underlying mechanisms of the fused cells resistant phenotype.

One likely explanation and our current hypothesis lies in the fused cell poly-aneuploidy state. Poly-aneuploid cells have shown general stress and therapeutic tolerance in many biological settings including a variety of cancers.^15–19^ Furthermore, poly-aneuploid human cells can have transcriptomic downregulation as many become senescent, and exit the cell cycle.^26^ More work is needed to be able to directly relate the fusion therapeutic resistance profile with their polyploid state, but the overall higher DNA content of fused cells demonstrated in **Figure 3** is a promising start.

Importantly, this work demonstrates a key insight into the potential clinical impacts of cell-cell fusion. While cell-cell fusion is a fundamentally rare event, the adaptive benefits it can provide seem to be selected for in tumor environments, particularly once treatments are administered. This idea is not completely novel; this approach to oncological research where tumor dynamics are evaluated through an evolutionary lens is known as evolutionary oncology.^7–9,35^ However, this perspective supports the further evaluation of cell-cell fusion as a future therapeutic target; rare events can still have tremendous clinical impacts.

Our recent review suggests that cell-cell fusion could become the next cancer hallmark in its own right, but that there is insufficient evidence for its occurrence and importance across cancer subtypes for this to be established at present.^2^ This study supports this claim by providing important initial identification and characterization of cell-cell fusion in NSCLC, and demonstrating its potential clinical impacts through a combination of drug sensitivity and transcriptomic profiling. We have demonstrated that the selective pressure of therapy can select for fitness advantages (e.g. therapeutic resistance) conferred following fusion. However, there are still many questions that need to be addressed before cell-cell fusion can be directly associated with rates of treatment failure, or targeted therapeutically. We have also shown the surprising differences in transcriptomic features between our three cell populations – parental, resistant and fused NSCLC cells – that in turn highlights how little we know about the underlying mechanisms and impacts of fusion on a cellular level. Understanding what contributes to the fused phenotypes and to what extent fused cells contribute to cancer progression and treatment failure remains to be seen, but the evidence in this paper suggests that there is a connection to explore more deeply.

## Methods

### Cell lines

Ancestral NSCLC PC-9 cells (Sigma-Aldrich, 90071810) were cultured in RPMI-1640 medium supplemented with 10% heat-inactivated fetal bovine serum and 1% penicillin/streptomycin. To create distinct drug-sensitive and drug-resistant populations, cells were labeled with eGFP and mCherry fluorescent proteins, respectively. Drug-sensitive cells were maintained in 0.1% DMSO as a vehicle control, while resistant cells were cultured in 1 µM gefitinib (Cayman, 13166) following protocols described by Farrokhian et al.^36^.

### Derivation of Fused Cells Using Co-culture experiments

To assess the long-term dynamics of cell-cell interactions and fitness across untreated and treated conditions, we co-cultured drug-sensitive and drug-resistant cell populations at five distinct ratios. Briefly, a total of 1 × 10^5^ cells, comprising both populations in varying proportions, were seeded into a 10cm^2^ tissue culture-treated dish and maintained in 0.1% DMSO as vehicle control for an initial period of six weeks. When cultures reached 80–90% confluence, cells were subcultured into fresh medium, and samples were collected at each subculturing interval to monitor population frequencies. These frequencies were analyzed using high-parameter, high-sensitivity flow cytometry (Sony ID70000). Following the initial six weeks, co-cultured populations were exposed to 20nM gefitinib and cultured under these conditions for an additional six weeks, with population frequencies similarly analyzed at each subculturing interval to assess proportions under drug selection pressure. After 12 weeks of co-culturing, double-positive cell populations that emerged were sorted and maintained in a medium containing 20nM gefitinib to study the stability and characteristics.

### Nuclear staining

To investigate nuclear fusion events in fused NSCLC cell populations compared to parental cell populations, we performed DAPI staining using 8-well chambered µ-slides (ibidi, 80806) followed by fluorescence microscopy. Cells were seeded at a density of 5 × 10^4^ per well. Once cells adhered to the slide, the medium was removed, and cells were fixed with 4% formaldehyde (prepared in 1X PBS) for 10 minutes at room temperature to preserve cellular structures. After fixation, formaldehyde was discarded, and samples were washed three times with 1X PBS. Cells were then permeabilized with a solution of 0.1% Triton X-100 in 1X PBS for 15 minutes at room temperature. Following permeabilization, the solution was removed, and a drop mounting media containing DAPI (Abcam, ab104139) was added to each well. Coverslips were carefully placed over the cells, and fluorescence images were captured using Leica fluorescence microscope to visualize nuclear fusion.

#### [article136||h3||]CELL Cycle Analysis

To assess cell cycle distribution, we utilized propidium iodide (PI) staining to quantify DNA content. Experimentally, cells were harvested and thoroughly washed with PBS to eliminate residual medium and drug. After washing, cells were fixed in cold 70% ethanol at 4°C for 30 minutes, then washed twice in PBS to remove the remaining ethanol. Samples were subsequently stained with FxCycle PI/RNase staining solution (Molecular Probes, F10797) following the manufacturer’s protocol. Stained samples were analyzed using flow cytometry, allowing for detailed determination of cell cycle distribution based on DNA content.

### Drug sensitivity profiling

To assess drug sensitivity, cells were harvested at 70–80% confluence and counted using 0.4% trypan blue solution (Thermo Fisher, 15250061) with Countess 3 Automated Cell Counter (Life Technologies). For cell viability assays, 3,000 cells per well in 100 µL of medium containing increasing concentrations of selected drugs were plated in 96-well microplates (Corning, 3904) in triplicate for drug-sensitive, drug-resistant, and fused cell populations. Plating was performed using a Multidrop reagent dispenser (Thermo Fisher) to ensure consistency. Drug-treated cells were incubated for 3 days, after which cell viability was measured using the CellTiter-Glo luminescent assay (Promega, G7571), with a luminescence signal read on a Tecan microplate reader. The resulting cell viability data were normalized to the average of DMSO-treated control and plotted as relative dose-response curves across the different cell types.

### Bulk RNAseq: Library Preparation, Quality Control, and Sequencing

Total RNA was extracted in triplicate from drug-sensitive PC-9, drug-resistant PC-9, and fused cell populations cultured as described above, using the Trizol RNA isolation reagent following the manufacturer’s instructions (Thermo Fisher Scientific, 15596026). RNA quality and integrity were evaluated using a NanoDrop spectrophotometer and Agilent TapeStation systems, while RNA concentration was determined using the Qubit RNA Broad Range (BR) assay kit (Thermo Fisher, Q10210). For library preparation, 500 ng of total RNA was used as input for the Illumina TruSeq RNA Library Prep Kit v2 (Illumina), following Illumina’s recommended protocol. The final libraries underwent quality assessment using TapeStation systems (Agilent) and quantification by Qubit (Thermo Fisher). Finally, sequencing was performed on the Illumina NovaSeq SP platform with 150 bp paired-end reads (2 × 150 bp).

### Bulk RNAseq: Sequence Alignment and Analyses

Read alignment and quantification is done via the STAR/Salmon workflow. We first filtered and adapter trimmed the paired-end reads using fastp with default settings. Subsequently the trimmed reads are aligned to GRCh38 primary assembly with gencode gene annotations v42 via STAR with two-pass mode and chimeric read detection enabled. Following the alignment, reads mapping to the transcript locations are quantified via Salmon with bootstrapping enabled. Transcript level quantifications (TPMs) are then used to calculate gene-level counts using the tximport package. Calculated gene-level counts are then used as input to edgeR based differential expression analysis workflow.

Following the transcript-level alignment, gene-level counts were generated using the tximport package^37^. Differential gene expression was run using the edgeR package^38^. Gene set enrichment (GSEA) and gene ontology enrichment analyses were performed using the clusterProfiler package.^39^ Plotting used a combination of ggplot2 and pheatmap.^40,41^

## Acknowledgements

JGS and PVS were supported by NIH 5R37CA244613-04 (https://www.cancer.gov/). PVS was supported by NIH 3T32GM007250-46S1 (https://www.nigms.nih.gov/). JGS was supported by American Cancer Society Research Scholar Grant RSG-20-096-01 (https://www.cancer.org/). DST was supported by The Research Council of Norway with grant 325628/ IAR (https://www.forskningsradet.no/en/). The funders had no role in study design, data collection and analysis, decision to publish, or preparation of the manuscript.

## Author contributions statement

PVS and DST contributed equally to this work. PVS designed and executed the majority of the analyses. DST was entirely responsible for *in vitro* experimental design and execution, with assistance from MND, SL, EAA, and SR. AD contributed extensively to the bioinformatics analyses and figure generation. Project conceptualization and groundwork was performed largely by RBC and DTW prior to PVS’s entrance to the lab. Manuscript writing and editing was done by PVS with extensive assistance from AD. Project was supervised by JGS.

---

**Supplemental Figure 1.**
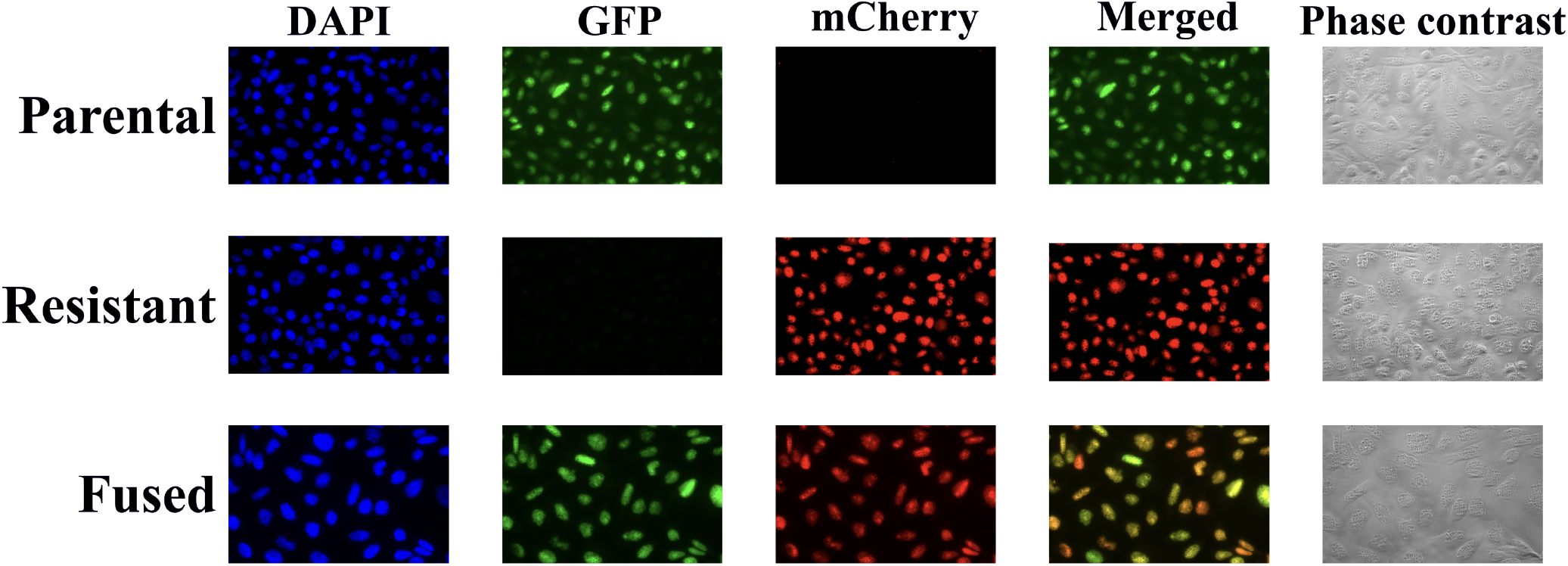
FACS images of parental, resistant, and fused cells.

**Supplemental Figure 2:**
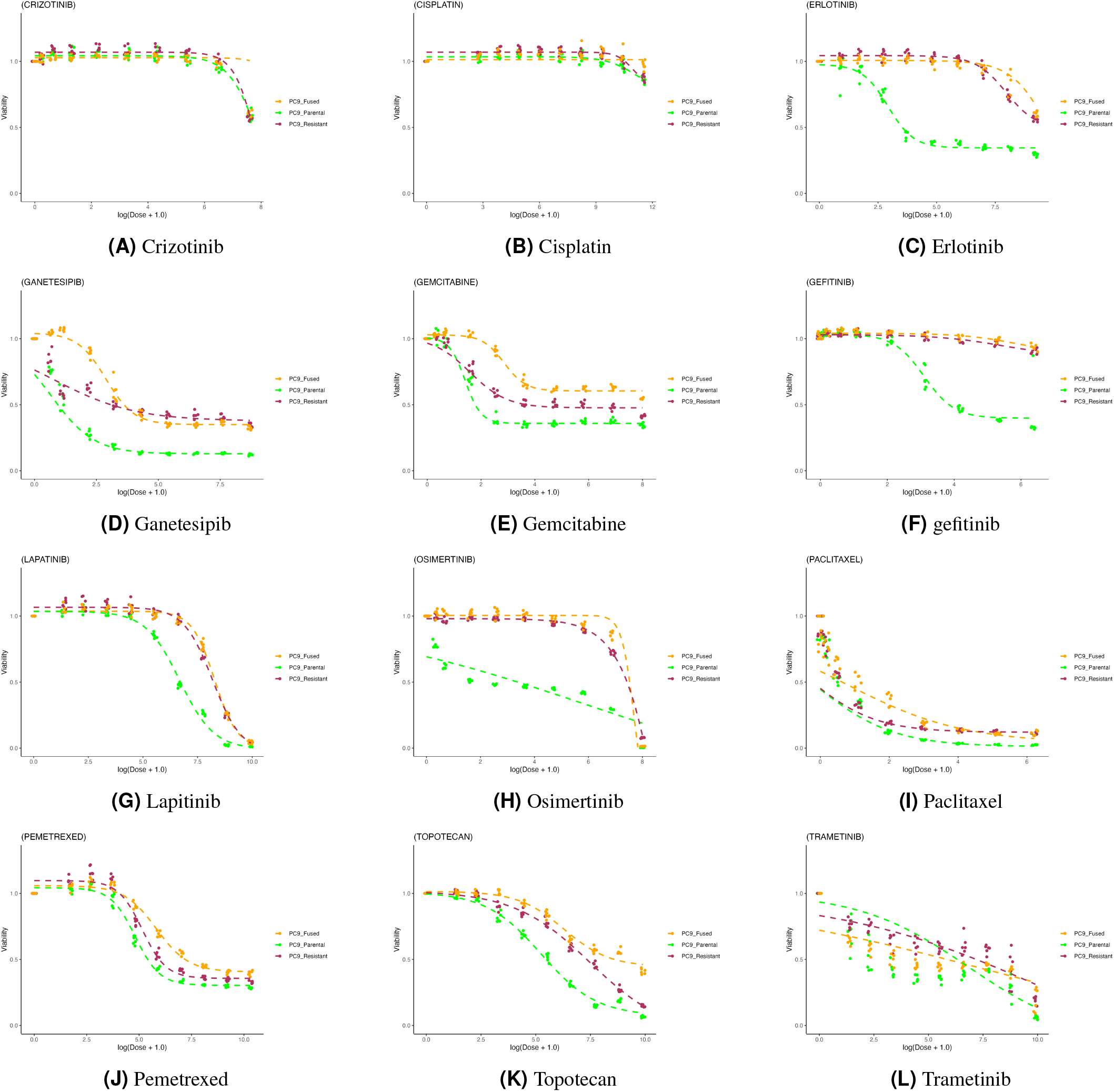
Selected fitted dose-response curves for fused, parental, and resistant cells across 12 drugs listed in alphabetical order.

**Supplemental Figure 3.**
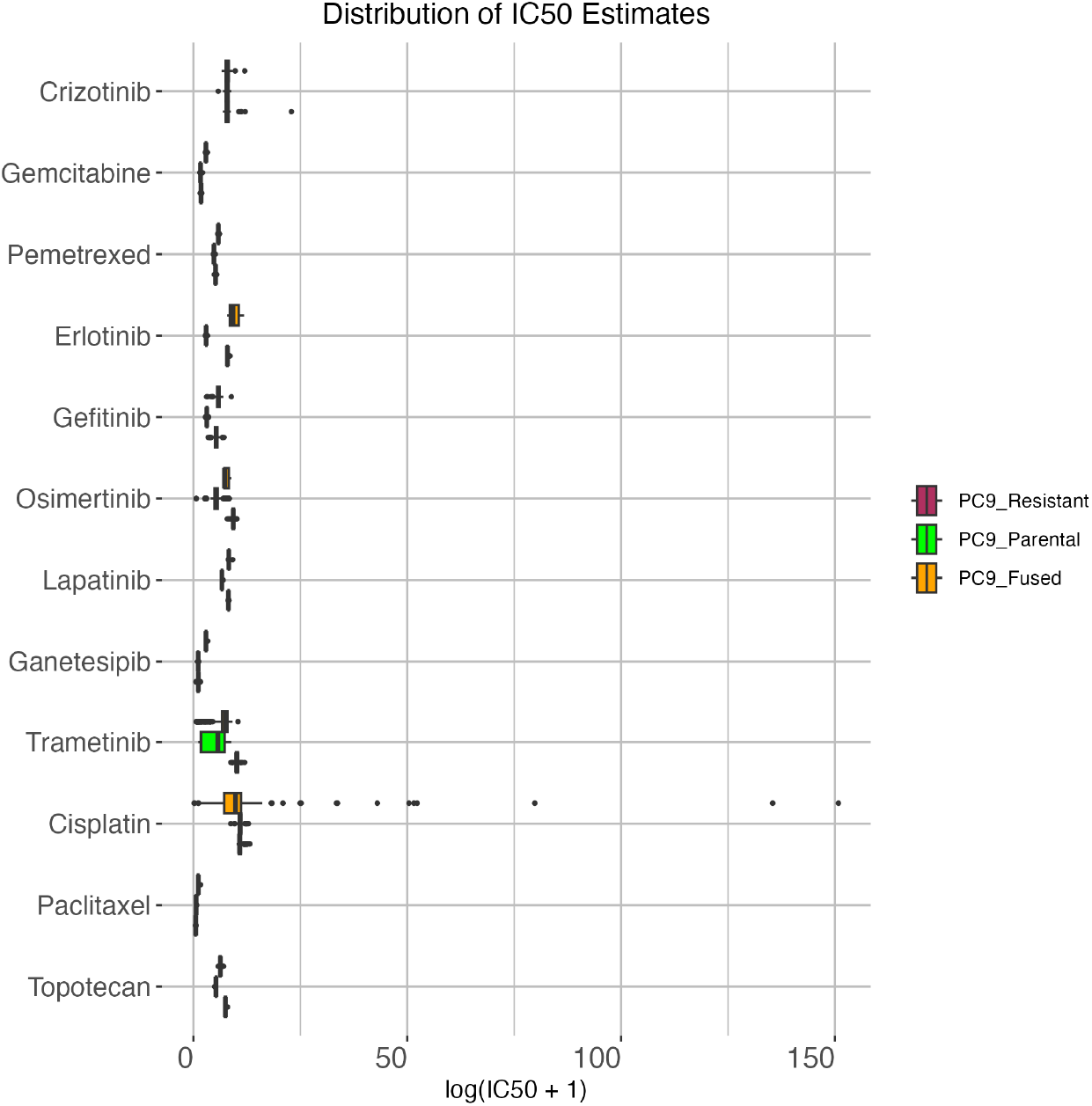
Extended Figure 4b. Repeated fitting of dose response curves (manual estimation of LL4 or hill function coefficients) leads to high variability in cisplatin response among fused cells.

## References

1. Hernández, J. M. & Podbilewicz, B. The hallmarks of cell-cell fusion, DOI: 10.1242/dev.155523 (2017). ISSN: 14779129 Issue: 24 Pages: 4481–4495 Publication Title: Development (Cambridge) Volume: 144.

2. Shultes, P. V., Weaver, D. T., Tadele, D. S., Barker-Clarke, R. J. & Scott, J. G. Cell-cell fusion in cancer: The next cancer hallmark? The Int. J. Biochem. Cell Biol. 175, 106649, DOI: 10.1016/j.biocel.2024.106649 (2024).

3. Brukman, N. G., Uygur, B., Podbilewicz, B. & Chernomordik, L. V. How cells fuse, DOI: 10.1083/jcb.201901017 (2019). ISSN: 15408140 Issue: 5 Pages: 1436-1451 Publication Title: Journal of Cell Biology Volume: 218.

4. Miroshnychenko, D. et al. Spontaneous cell fusions as a mechanism of parasexual recombination in tumour cell populations. Nat. Ecol. Evol. 5, 379–391, DOI: 10.1038/s41559-020-01367-y (2021). Publisher: Nature Research.

5. Hendriks, L. E. L. et al. Non-small-cell lung cancer. Nat. Rev. Dis. Primers 10, 71, DOI: 10.1038/s41572-024-00551-9 (2024).

6. Yeldag, G., Rice, A. & del Río Hernández, A. Chemoresistance and the Self-Maintaining Tumor Microenvironment. Cancers 10, 471, DOI: 10.3390/cancers10120471 (2018).

7. Scarborough, J. A., Eschrich, S. A., Torres-Roca, J., Dhawan, A. & Scott, J. G. Exploiting convergent phenotypes to derive a pan-cancer cisplatin response gene expression signature. NPJ Precis. Oncol. 7, 38 (2023).

8. King, E. S., Tadele, D. S., Pierce, B., Hinczewski, M. & Scott, J. G. Diverse mutant selection windows shape spatial heterogeneity in evolving populations. PLOS Comput. Biol. 20, 1–22, DOI: 10.1371/journal.pcbi.1011878 (2024).

9. Maltas, J. et al. Frequency-dependent ecological interactions increase the prevalence, and shape the distribution, of preexisting drug resistance. PRX Life 2, 023010, DOI: 10.1103/PRXLife.2.023010 (2024).

10. Tajima, Y., Shibasaki, F. & Masai, H. Cell fusion upregulates PD-L1 expression for evasion from immunosurveillance. Cancer Gene Ther. 31, 158–173, DOI: 10.1038/s41417-023-00693-0 (2024).

11. Gluzman, M., Scott, J. G. & Vladimirsky, A. Optimizing adaptive cancer therapy: dynamic programming and evolutionary game theory. Proc. Royal Soc. B 287, 20192454 (2020).

12. Basanta, D. et al. Investigating prostate cancer tumour–stroma interactions: clinical and biological insights from an evolutionary game. Br. journal cancer 106, 174–181 (2012).

13. Wölfl, B. et al. The contribution of evolutionary game theory to understanding and treating cancer. Dyn. Games Appl. 12, 313–342 (2022).

14. Hass, R., Ohe, J. v. d. & Dittmar, T. Hybrid formation and fusion of cancer cells in vitro and in vivo, DOI: 10.3390/cancers13174496 (2021). ISSN: 20726694 Issue: 17 Publication Title: Cancers Volume: 13.

15. Tong, K. et al. Genome duplication in a long-term multicellularity evolution experiment. Nature 1–9 (2025).

16. Addissouky, T. A. Polyploidy-mediated resilience in hepatic aging: molecular mechanisms and functional implication. Egypt. Liver J. 14, 83 (2024).

17. Coward, J. & Harding, A. Size does matter: why polyploid tumor cells are critical drug targets in the war on cancer. Front. oncology 4, 123 (2014).

18. Schmidt, M. J. et al. Polyploid cancer cells reveal signatures of chemotherapy resistance. Oncogene 44, 439–449 (2025).

19. Song, Y., Zhao, Y., Deng, Z., Zhao, R. & Huang, Q. Stress-Induced Polyploid Giant Cancer Cells: Unique Way of Formation and Non-Negligible Characteristics, DOI: 10.3389/fonc.2021.724781 (2021). ISSN: 2234943X Publication Title: Frontiers in Oncology Volume: 11.

20. Alduais, Y., Zhang, H., Fan, F., Chen, J. & Chen, B. Non-small cell lung cancer (NSCLC): A review of risk factors, diagnosis, and treatment. Medicine 102 (2023).

21. Guo, Q. et al. Current treatments for non-small cell lung cancer. Front. Oncol. 12, DOI: 10.3389/fonc.2022.945102 (2022).

22. Zhang, S. et al. Generation of Cancer Stem-like Cells through Formation of Polyploid Giant Cancer Cells. Oncogene 33, 10.1038/onc.2013.96, DOI: 10.1038/onc.2013.96 (2014).

23. Uygur, B. et al. Interactions with muscle cells boost fusion, stemness, and drug resistance of prostate cancer cells. Mol. Cancer Res. 17, 806–820, DOI: 10.1158/1541-7786.MCR-18-0500 (2019). Publisher: American Association for Cancer Research Inc.

24. Dittmar, T. Generation of Cancer Stem/Initiating Cells by Cell–Cell Fusion, DOI: 10.3390/ijms23094514 (2022). ISSN: 14220067 Issue: 9 Publication Title: International Journal of Molecular Sciences Volume: 23.

25. Melzer, C., Ohe, J. v. d. & Hass, R. In Vitro Fusion of Normal and Neoplastic Breast Epithelial Cells with Human Mesenchymal Stroma/Stem Cells Partially Involves Tumor Necrosis Factor Receptor Signaling. Stem Cells 36, 977–989, DOI: 10.1002/stem.2819 (2018). Publisher: Wiley-Blackwell.

26. Hass, R., von der Ohe, J. & Dittmar, T. Cancer Cell Fusion and Post-Hybrid Selection Process (PHSP). Cancers 13, 4636, DOI: 10.3390/cancers13184636 (2021). Number: 18 Publisher: Multidisciplinary Digital Publishing Institute.

27. Dietz, M. S. et al. Relevance of circulating hybrid cells as a non-invasive biomarker for myriad solid tumors. Sci. Reports 11, 1–13, DOI: 10.1038/s41598-021-93053-7 (2021). Number: 1 Publisher: Nature Publishing Group.

28. Bateman, A., Bullough, F. & Murphy, S. Fusogenic Membrane Glycoproteins As a Novel Class of Genes for the Local and Immune-mediated Control of Tumor Growth 1 (2000). Pages: 1492-1497 Publication Title: CANCER RESEARCH Volume: 60.

29. Tretyakova, M. S., Subbalakshmi, A. R., Menyailo, M. E., Jolly, M. K. & Denisov, E. V. Tumor Hybrid Cells: Nature and Biological Significance. Front. Cell Dev. Biol. 10 (2022).

30. Shin, J. H. et al. Colon cancer cells acquire immune regulatory molecules from tumor-infiltrating lymphocytes by trogocytosis. Proc. Natl. Acad. Sci. 118, e2110241118, DOI: 10.1073/pnas.2110241118 (2021). Publisher: Proceedings of the National Academy of Sciences.

31. Dörnen, J., Myklebost, O. & Dittmar, T. Cell fusion of mesenchymal stem/stromal cells and breast cancer cells leads to the formation of hybrid cells exhibiting diverse and individual (stem cell) characteristics. Int. J. Mol. Sci. 21, 9636 (2020).

32. Bühler, A. et al. When Mechanical Stress Matters: Generation of Polyploid Giant Cancer Cells in Tumor-like Microcapsules, DOI: 10.1101/2022.09.22.508846 (2022). Pages: 2022.09.22.508846 Section: New Results.

33. Demin, S., Berdieva, M. & Goodkov, A. Cell-cell fusions and cell-in-cell phenomena in healthy cells and cancer: Lessons from protists and invertebrates. Semin. Cancer Biol. 81, 96–105, DOI: 10.1016/j.semcancer.2021.03.005 (2022).

34. Dittmar, T., Weiler, J., Luo, T. & Hass, R. Cell-Cell Fusion Mediated by Viruses and HERV-Derived Fusogens in Cancer Initiation and Progression. Cancers 13, 5363, DOI: 10.3390/cancers13215363 (2021). Number: 21 Publisher: Multidisciplinary Digital Publishing Institute.

35. Cairns, J. Mutation selection and the natural history of cancer. Nature 255, 197–200, DOI: 10.1038/255197a0 (1975).

36. Farrokhian, N. et al. Measuring competitive exclusion in non–small cell lung cancer. Sci. Adv. 8, eabm7212 (2022).

37. Soneson, C., Love, M. I. & Robinson, M. D. Differential analyses for rna-seq: transcript-level estimates improve gene-level inferences. F1000Research 4, DOI: 10.12688/f1000research.7563.1 (2015).

38. Robinson, M. D., McCarthy, D. J. & Smyth, G. K. edger: a bioconductor package for differential expression analysis of digital gene expression data. Bioinformatics 26, 139–140, DOI: 10.1093/bioinformatics/btp616 (2010).

39. Yu, G., Wang, L.-G., Han, Y. & He, Q.-Y. clusterprofiler: an r package for comparing biological themes among gene clusters. OMICS: A J. Integr. Biol. 16, 284–287, DOI: 10.1089/omi.2011.0118 (2012).

40. Wickham, H. ggplot2: Elegant Graphics for Data Analysis (Springer-Verlag New York, 2016).

41. Kolde, R. pheatmap: Pretty Heatmaps (2019). R package version 1.0.12.

